# Aversion encoded in the subthalamic nucleus

**DOI:** 10.1101/2020.07.09.195610

**Authors:** Gian Pietro Serra, Adriane Guillaumin, Jérome Baufreton, François Georges, Åsa Wallén-Mackenzie

**Affiliations:** Department of Organism Biology, Uppsala University, SE-752 36 Uppsala, Sweden; Université de Bordeaux, Institut des Maladies Neurodégénératives, UMR 5293, F-33000 Bordeaux, France; CNRS, Institut des Maladies Neurodégénératives, UMR 5293, F-33000 Bordeaux, France

## Abstract

Activation of the subthalamic nucleus (STN) is associated with the stopping of ongoing behavior via the basal ganglia. However, we recently observed that optogenetic STN excitation induced a strong jumping/escaping behavior. We hypothesized that STN activation is aversive. To test this, place preference was assessed. Optogenetic excitation of the STN caused potent place aversion. Causality between STN activation and aversion has not been demonstrated previously. The lateral habenula (LHb) is a critical hub for aversion. Optogenetic stimulation of the STN indeed caused firing of LHb neurons, but with delay, suggesting the involvement of a polysynaptic circuit. To unravel a putative pathway, the ventral pallidum (VP) was investigated. VP receives projections from the STN and in turn projects to the LHb. Optogenetic excitation of STN-VP terminals caused firing of VP neurons and induced aversive behavior. This study identifies the STN as critical hub for aversion, potentially mediated via an STN-VP-LHb pathway.

## Introduction

The subthalamic nucleus (STN) is a small and bilaterally positioned nucleus which exerts excitatory influence over the indirect pathway of the basal ganglia (Kita, 1994; Kita and Kitai, 1987). The STN has long been associated with motor and cognitive processes (Bonnevie and Zaghloul, 2019; Schmidt and Berke, 2017; Weintraub and Zaghloul, 2013). Aberrant STN activity is observed in patients with Parkinson’s disease and obsessive-compulsive disorder (OCD), but can be corrected by high-frequency electrical stimulation of the STN, so called deep brain stimulation (DBS), which successfully alleviates both motor deficiency in Parkinson’s disease and compulsivity in OCD (Benabid et al., 2009; Blomstedt et al., 2015; Mallet et al., 2008). However, while STN-DBS is highly beneficial to many severely afflicted individuals, a number of adverse side-effects have been reported, including low emotional state (Péron et al., 2013; Temel et al., 2006). This has not been easily understood, but highlights the need for revealing the full repertoire of regulatory roles executed by the STN.

Reward and punishment motivate learning through complex neuronal circuits (Jean-Richard-Dit-Bressel et al., 2018). Studies of rewarding and aversive stimuli were long focused on dopamine neurons in the ventral tegmental area (VTA) (Barker et al., 2016; Bromberg-Martin et al., 2010; Lammel et al., 2014; Wise, 2004). However, with the seminal discovery that glutamatergic neurons of the lateral habenula (LHb) respond to reward in an opposite manner as dopamine neurons, LHb has gained considerable attention as an important brain area in the regulation of aversive responses and negative reward-processing (Matsumoto and Hikosaka, 2007). LHb neurons are activated both by aversive stimuli and the omission of an expected reward (Gao et al., 1996; Matsumoto and Hikosaka, 2007). Further, LHb hyperactivity has been associated with depression and negative symptoms of addiction (Hu, 2016).

The LHb receives regulatory input from several sources, many identified over the past few years using optogenetic stimulations. For example, optogenetic excitation of glutamatergic projections from the VTA and the lateral hypothalamus was shown to induce aversion (Lazaridis et al., 2019; Lecca et al., 2017; Root et al., 2014; Stamatakis et al., 2016), as did also stimulation of afferent fibers from the entopenduncular nucleus (EP) (Shabel et al., 2012; Stephenson-Jones et al., 2016). In addition, optogenetic stimulation of glutamatergic neurons in the ventral pallidum (VP) was recently shown to evoke post-synaptic activity in the LHb and to drive place avoidance (Faget et al., 2018; Tooley et al., 2018). Thus, through various sources of excitatory input, activation of the LHb mediates aversive behavior.

The STN projects directly to several of the structures recognized in aversive processing, including the EP and VP, both directly connected to the LHb (Faget et al., 2018; Shabel et al., 2012; Stephenson-Jones et al., 2016; Tooley et al., 2018). However, while ample studies have addressed pallidal and habenular structures using optogenetics leading towards increased understanding of their role in affective functions, only a limited set of optogenetic studies have yet investigated the STN. Based on the strong interest in STN-DBS, optogenetics in the STN has mainly been used to address motor functions in the context of Parkinson’s disease (Gradinaru et al., 2009; Sanders and Jaeger, 2016; Tian et al., 2018; Yoon et al., 2016, 2014). Further, one recent study identified that optogenetic excitation of the STN interrupted ongoing licking behavior in mice (Fife et al., 2017). STN neurons express the *Pitx2* gene, which allows recombination of floxed alleles using Pitx2-Cre transgenic mice (Martin et al., 2004). Confirming the selectivity of the Pitx2-Cre transgene in directing opsin expression to the STN, we have previously shown that photostimulation of the STN results in post-synaptic currents and glutamate release in basal ganglia target areas (Schweizer et al., 2016, 2014; Viereckel et al., 2018). In a recent report, we show that optogenetic excitation and inhibition result in opposite locomotor effects (Guillaumin et al., 2020). Surprisingly, we also observed that STN excitation could induce an avoidance-like behavior. When placed in an open and novel arena, mice displayed a strong escaping behavior, characterized by attempts to jump out of the apparatus. This unexpected outcome was evident immediately upon STN excitation (Guillaumin et al., 2020).

Based on these observations, we hypothesized that the STN is engaged in aversive behavior. To test this, optogenetic stimulations followed through with electrophysiological recordings and place preference analyses were implemented in Pitx2-Cre mice. Our results identify place avoidance upon selective activation of either the STN itself or STN terminals in the VP, and also demonstrate multi-synaptic STN-communication with the LHb. The findings contribute to the current decoding of the neurocircuitry of aversive behavior.

## Results

### STN neurons follow 20 Hz optogenetic stimulation

Optogenetics allows spatially and temporally precise control of neuronal activation which under experimental conditions enables the correlation between distinct neuronal activation and measurable behaviors in freely-moving animals (Deisseroth, 2011). To allow optogenetic control of STN neurons, Pitx2-Cre mice were bilaterally injected with adeno-associated (AAV) virus to express a construct encoding either Channelrhodopsin-2 (ChR2) fused with eYFP (so called Pitx2/ChR2-eYFP mice) or, as controls, eYFP alone (Pitx2/eYFP; control mice) (Figure 1A). Bilateral optic cannulas were placed above the STN (Figure 1A). Histological analyses and *in vivo* extracellular recordings were performed to validate the approach, and injections and cannulas placement were confirmed after completed behavioral analyses.

**Figure 1.**
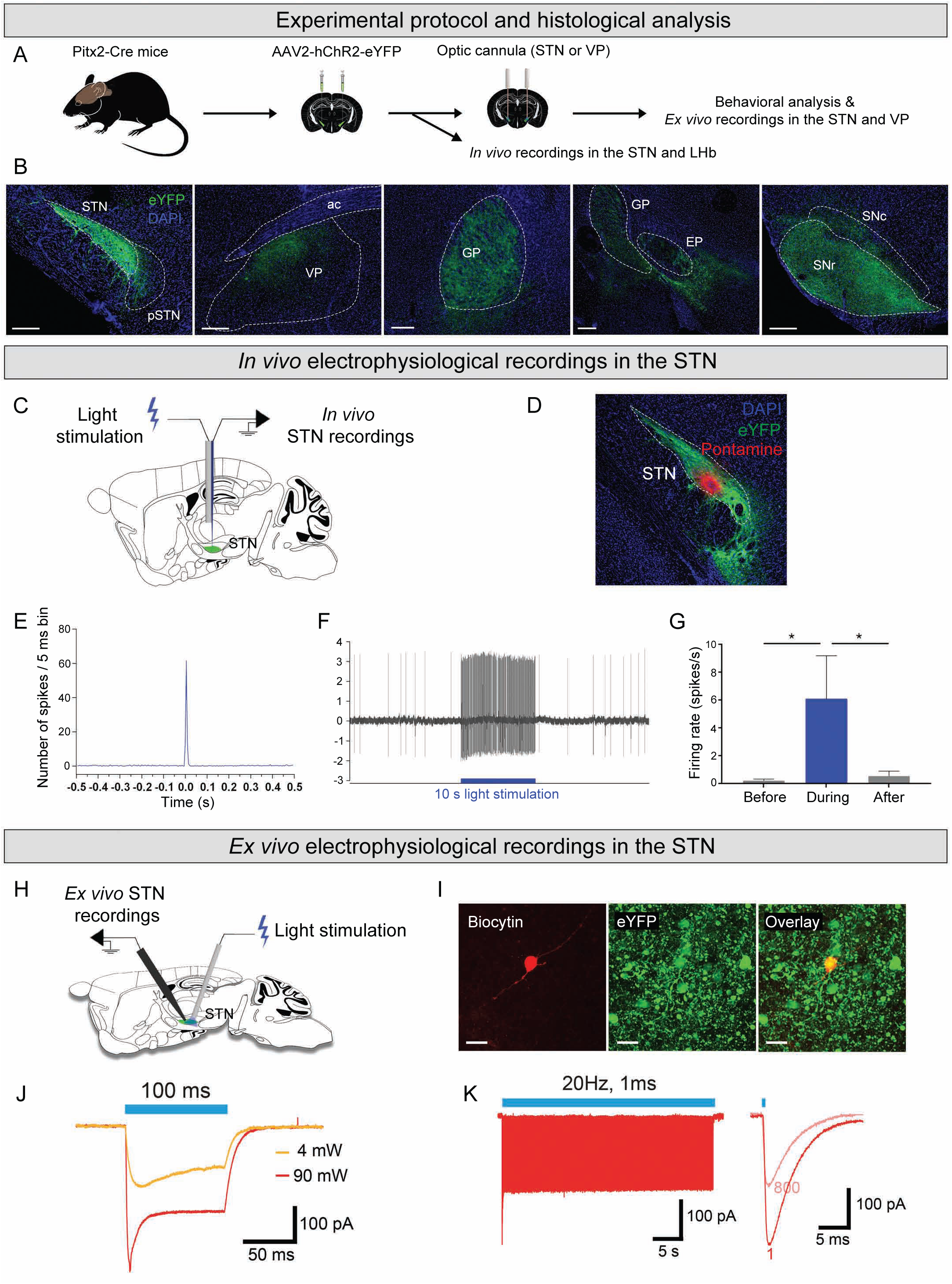
Photostimulation in the STN in Pitx2-ChR2-eYFP mice excites STN neurons. (A) Graphical representation of stereotaxic bilateral injection into the STN of an rAAV carrying a Cre-dependent Channelrhodopsin 2 (ChR2) fused with eYFP (ChR2-eYFP), or eYFP alone, into Pitx2-Cre mice (Pitx2/ChR2-eYFP mice and Pitx2/eYFP control mice, respectively) followed by canula implantation above the STN. (B) Coronal sections showing eYFP fluorescence (green) in STN neurons and its target areas projections in a representative Pitx2/ChR2-eYFP mouse; from left to right: STN cell bodies, STN target areas projections in the VP, GP, EP, SNr and SNc; nuclear marker (DAPI, blue); scale, 250 µm. (C) Procedure for STN optotagging experiments. (D) eYFP-positive neurons (green) in the STN; pontamine blue sky deposit (red dot) at the last recorded coordinate; nuclear marker (DAPI, blue) (scale, 200 µm). (E) Averaged PSTH of STN-excited neurons upon light stimulation (0.5 Hz, 5 ms pulse duration, 5-8 mW). (F) Example of an excited STN neuron during a behavioral protocol (20 Hz, 5 ms pulse duration, 5-8 mW) with the frequency and the recording trace. (G) Frequency of STN neurons before, during and after 10 s light stimulation following the behavioral protocol (20 Hz, 5 ms pulse duration, 5-8 mW); data expressed as mean ± SEM, n=7, both *p=0.15. (H) Graphical representation of a para-sagittal slice obtained from a Pitx2/ChR2-eYFP mouse with a patch-clamp electrode and an optic fiber placed above the STN. (I) Confocal images depicting a STN neuron filled with biocytin (BC) and expressing the ChR2-eYFP construct (visualized by eYFP). Scale bar: 10µm. (J) Representative example of typical ChR2-mediated currents induced by a 1 s blue light (λ = 470 nm) pulses at intensities of 4mW and 90 mW, respectively. (K) ChR2-mediated currents induced by a train of blue light of 40 s duration at 20 Hz (individual light pulses of 1 ms). Abbreviations: STN, subthalamic nucleus; VP, ventral pallidum; GP, globus pallidus, EP; entopenduncular nucleus; SNr, substantia nigra *pars reticulata*; SNc, substantia nigra *pars compacta*.

Mice that displayed strong cellular eYFP labeling throughout the extent of the STN and in which optic cannulas could be confirmed as positioned immediately above the STN, were included in the statistical analyses of the electrophysiological and behavioral experiments (Figure 1B). In addition to the STN, the para-subthalamic nucleus (pSTN), associated with the medial STN, was eYFP-labeled to minor extent. Addressing eYFP-positive terminals in target areas of the STN, these were detected in the GP (GP externa in humans), EP, substantia nigra *pars reticulata* (SNr), substantia nigra *pars compacta* (SNc) and VP, confirming that Pitx2-Cre-positive neurons of the STN reach these target areas (Figure 1B).

To ensure connectivity, *in vivo* single cell electrophysiological recordings upon optogenetic stimulation of the STN were performed (Figure 1C,D). First, an optotagging protocol (Figure 1E) was used to stimulate and record within the STN. To observe the reaction of STN neurons to photostimulation, peri-stimulus time histograms (PSTH protocol, 0.5 Hz, 5 ms bin width, 5-8 mW) were created by applying a 0.5 Hz stimulation protocol for at least 100 seconds. Action potentials in ChR2-positive STN cells were successfully evoked by STN photostimulation (Figure 1E).

Once neuronal activity of STN neurons returned to baseline, a photostimulation protocol intended for behavioral experiments was validated (Behavior protocol, 20 Hz, 5 ms pulses, 5-8 mW; 10 seconds) which increased the frequency and firing rate of STN neurons for the whole duration of the stimulation, after which they returned to normal (Figure 1F).

The excitability of STN neurons was next confirmed by patch-clamp recordings in the STN of Pitx2/ChR2-eYFP mice with the optic fiber placed above the recording site (Figure 1G). ChR2-eYFP expression was strong in the STN (Figure 1H), and all of the STN neurons tested responded to continuous (Figure 1I) or 20 Hz trains (Figure 1J) of light stimulation by sustained ChR2-mediated currents. When light stimulation was applied in brain slices from Pitx2/eYFP, no current were observed in STN neurons (n=3, data not shown).

Having confirmed optogenetics-driven activation of STN neurons, behavioral effects upon optogenetic STN activation were next tested.

### Activation of STN neurons induces place avoidance

To experimentally test the hypothesis that the STN drives affective behavior, we applied optogenetic excitation in the real-time place preference paradigm (RT-PP) and the elevated plus maze (EPM), representing validated tests for aversion and anxiety. First, Pitx2/ChR2-eYFP and control mice were analyzed in the RT-PP. Here, the mouse moves around freely in a three-compartment apparatus, in which entry into one of two main compartments (A or B) is paired with photostimulation of the STN. A photostimulation-neutral, transparent, corridor (neutral compartment) connects the main compartments, allowing free access to the whole apparatus. In this setup, an aversive response to photostimulation would be displayed as avoidance of entry into the light-paired compartment, and hence avoidance of activation of the STN. Further, if the association is strong, this would be expected to give rise to similar behavior even during days after the experience of STN activation due to conditioning effect (Figure 2A). Pitx2/ChR2-eYFP mice showed an active avoidance of STN-photostimulation by spending significantly less time in the light-paired compartment (A) than the unpaired compartment (B) both during the two days of real-time exposure to photostimulation (Days 3, 4), and also during the conditioned test days (Days 5, 6) (Figure 2B,C). Control mice showed an absence of stimulation-dependent response throughout the trials (Figure 2B).

**Figure 2.**
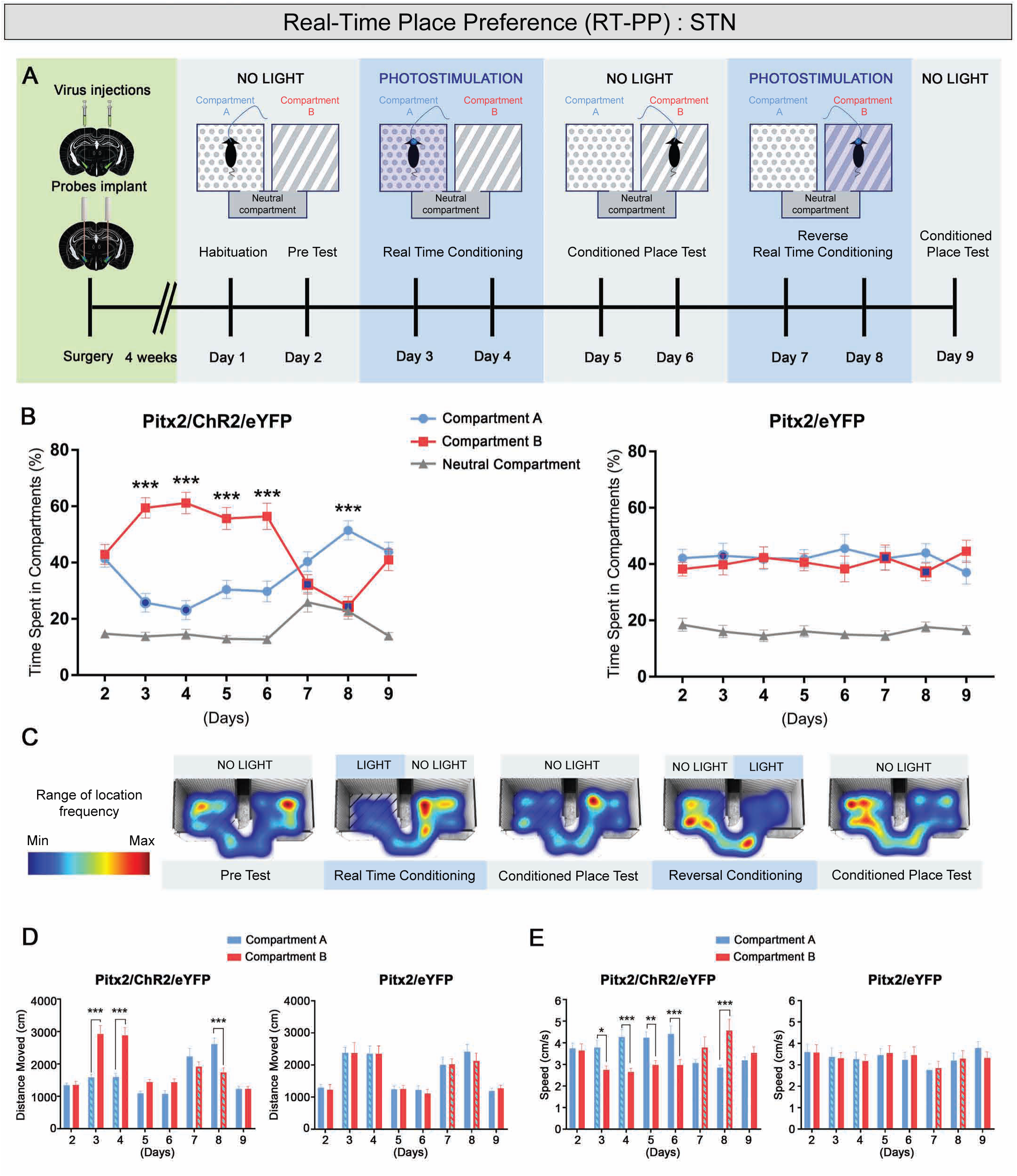
Optogenetic activation of STN neurons induces place aversion. (A) Graphical representation of the real-time place preference (RT-PP) setup in which Pitx2/ChR2-eYFP and control mice were tested. (B) Percentage of time that Pitx2/ChR2-eYFP and control mice spent in each compartment express as mean ± SEM. Dark blue filled circles (compartment A) and dark blue filled squares (compartment B) indicate photostimulation in that compartment. (Left panel) n=12 Pitx2/ChR2-eYFP, compartment A vs. compartment B, ***p<0.001; (Right panel) n=10 Pitx2/eYFP controls. (C) Representative occupancy heat-maps for Pitx2/ChR2-eYFP mice show less time spent in the photostimulation-paired chamber than in the unpaired chamber. (D) Distance moved in each compartment expressed as mean ± SEM throughout the course of the experiment. Light blue filled histograms indicate photostimulation in that compartment. (Left panel), ***p<0.001. (Right panel) n=10 Pitx2/eYFP control mice. (E) Speed in each compartment expressed as mean ± SEM. (Left panel) n=12 Pitx2/ChR2-eYFP mice, *p<0.05, **p<0.01, ***p<0.001. (Right panel) n=10 Pitx2/eYFP control mice.

Next, to test the ability for reversal of the avoidance behavior, a strategy was implemented in which the previously light-unpaired compartment was paired with photostimulation, and the stimulation protocol was repeated. On the first reversal day (Day 7), Pitx2/ChR2-eYFP mice responded by spending time in all three compartments which resulted in lack of significant preference for any compartment. On the second day (Day 8), mice again spent significantly more time in the unpaired compartment (A). On the final conditioned test day (Day 9), Pitx2/ChR2-eYFP mice spent a similar amount of time in the main compartments A and B (Figure 2B,C).

Avoidance towards the light-paired compartment was corroborated by faster and shorter distance moved compared to the non-light paired compartment across all conditioning days, except Day 7 when no specific compartment was preferred or avoided (Figure 2D,E). Control mice spent a comparable amount of time in A and B regardless of which compartment the stimulation was applied in, and showed no significant effects in distance moved or speed (Figure 2D,E). These findings identified a direct correlation between STN activation and place avoidance both upon direct STN stimulation and upon exposure to aversion-paired cues.

### Aversion induced by optogenetic STN activation stronger than inherent aversion for unsheltered exposure

To assess a putative anxiogenic component and to further pinpoint the nature of the aversive response observed in the RT-PP, mice were next analyzed in two versions of the EPM, the regular EPM and a modified EPM setup established to compare any aversion to STN-stimulation with a naturally occurring aversive situation (Figure 3A,D).

**Figure 3.**
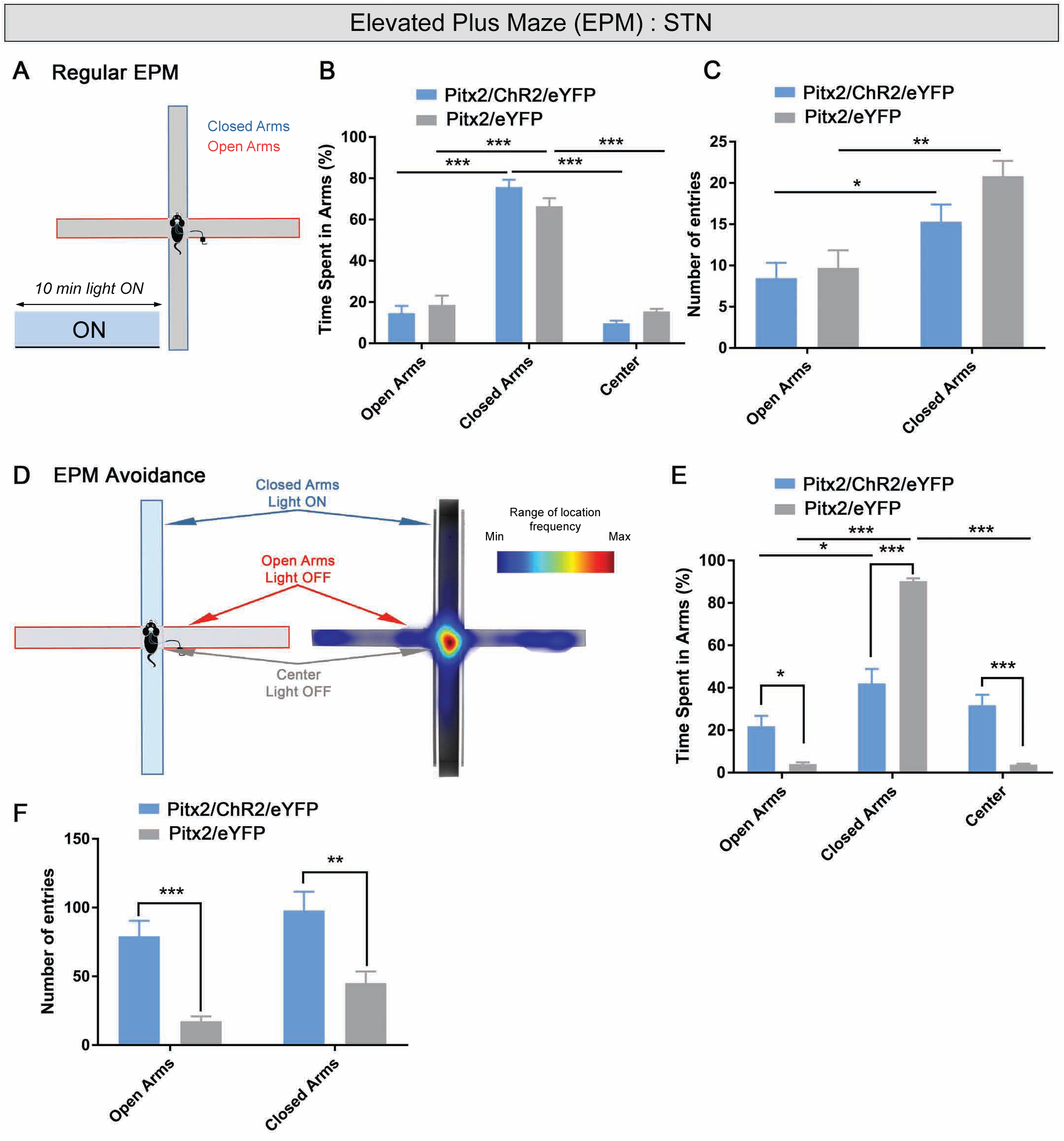
Aversion induced by optogenetic STN activation stronger than inherent aversion for unsheltered exposure. (A) Schematic representation of the elevated plus maze (EPM) in which Pitx2/ChR2-eYFP and control mice were tested; STN-photostimulation delivered for 10 minutes throughout the entire test arena. (B) Percentage of time spent in arms expressed as mean ± SEM, n=17 Pitx2/ChR2/eYFP, open arms vs. closed arms ***p<0.001; center vs. closed arms, ***p<0.001; n=11 Pitx2/eYFP mice, open arms vs. closed arms, ***p<0.001; center vs. closed arms, ***p<0.001. (C) Number of entries in arms expressed as mean ±SEM, n=17 Pitx2/ChR2/eYFP mice, open arms vs. closed arms *p< 0.05; n=11 Pitx2/eYFP, open arms vs. closed arms, **p< 0.01. (D) Graphical representation of a modified version of the EPM test and representative heat map showing occupancy. (E) Time spent in arms expressed as mean ±SEM, n=14 Pitx2/ChR2/eYFP, open arms vs. closed arms, *p<0.05; n=11 Pitx2/eYFP, open arms vs. closed arms, ***p<0.001; center vs. closed arms, ***p<0.001; Pitx2/ChR2/eYFP vs. Pitx2/eYFP, open arms *p<0.05, closed arms ***p<0.001, center ***p<0.001. (F) Number of entries in arms expressed as mean ±SEM, Pitx2/ChR2/eYFP vs. Pitx2/eYFP, open arms ***p<0.001; Closed arms **p<0.01.

The regular EPM is commonly used to assess anxiety. Here, the innate response of mice is their aversion towards the exposed open arms and a center zone, resulting in an avoidance of these areas, and consequently, a preference for the sheltered space provided by the closed arms. In the current experiment, STN-photostimulation was given throughout the trial (Figure 3A). Similar as controls, Pitx2/ChR2-eYFP mice spent most of the time in the sheltered space provided by the closed arms where duration was significantly higher compared to exposed open arms and the center (Figure 3B). Number of entries into either the open or closed arms was not significantly different between Pitx2/ChR2-eYFP and control mice, but for both groups, the number of entries into the closed arms was statistically higher than entry into open arms (Figure 3C). Since Pitx2/ChR2-eYFP mice showed the same behavior as control mice, the results suggest that activation of the STN is not associated with altered response towards the naturally aversive area, which they actively avoid both in presence and absence of STN stimulation. These results suggest that the aversive behavior upon STN-activation is likely not related to an anxiogenic effect.

Next, we re-designed the EPM test to directly compare the optogenetically induced aversive behavior with the naturally aversive context offered by the exposed open arm and center zone. In this EPM-avoidance setup, STN-photostimulation was activated directly upon entry into one of the two closed arms and disabled by leaving them, while entry into open arms or the center zone had no effect on STN-photostimulation (Figure 3D). The effect was striking; Pitx2/ChR2-eYFP mice actively tried to avoid the previously preferred closed arms, showed increased number of entries in all arms, and spent significantly more time in the open arms and in the center (Figure 3E,F). These findings demonstrated that the aversive effect induced by STN activation was stronger than the naturally aversive effect provided by unsheltered exposure. Thus, while not confirming STN activation as anxiogenic, analysis in both the RT-PP and EPM-avoidance paradigms demonstrated a causal relationship between STN activation and aversive behavior.

### STN photoactivation induces excitatory post-synaptic responses in the LHb neurons revealing poly-synaptic connectivity

The LHb has recently emerged as an important brain area for aversive behavior, firmly demonstrated by optogenetic analyses of the LHb itself as well as projections to the LHb (Faget et al., 2018; Lazaridis et al., 2019; Lecca et al., 2017; Shabel et al., 2012; Stephenson-Jones et al., 2016; Tooley et al., 2018). While STN activation has not previously been associated with aversion, it has been shown that high frequency stimulation of the STN in rats induces c-Fos expression in LHb (Tan et al., 2011) and modulation of LHb neuron activity (Hartung et al., 2016).

The potential impact of STN-photostimulation on LHb activity was assessed using *in vivo* extracellular recordings in Pitx2/ChR2-eYFP (Figure 4A). An optic fiber was positioned above the STN (Figure 4C), and a stimulation protocol (PSTH 0.5 Hz, 5 ms pulses, 5-8 mW; at least 100 seconds) was implemented which evoked an excitatory response in 50% of the recorded LHb neurons (onset latency =10.08 ms +/− 0.81 ms) (Figure 4B). Pontamine staining and neurobiotin-marked neurons confirmed the positioning of the recording electrodes within the LHb and enabled the reconstruction of neuronal positions (Figure 4D,E). No eYFP-positive fibers were detected in the habenular area of Pitx2/ChR2-eYFP (Figure 4E), confirming the absence of a direct connection between the STN and the LHb. These findings indicate a multi-synaptic glutamatergic pathway between the STN and the LHb.

**Figure 4.**
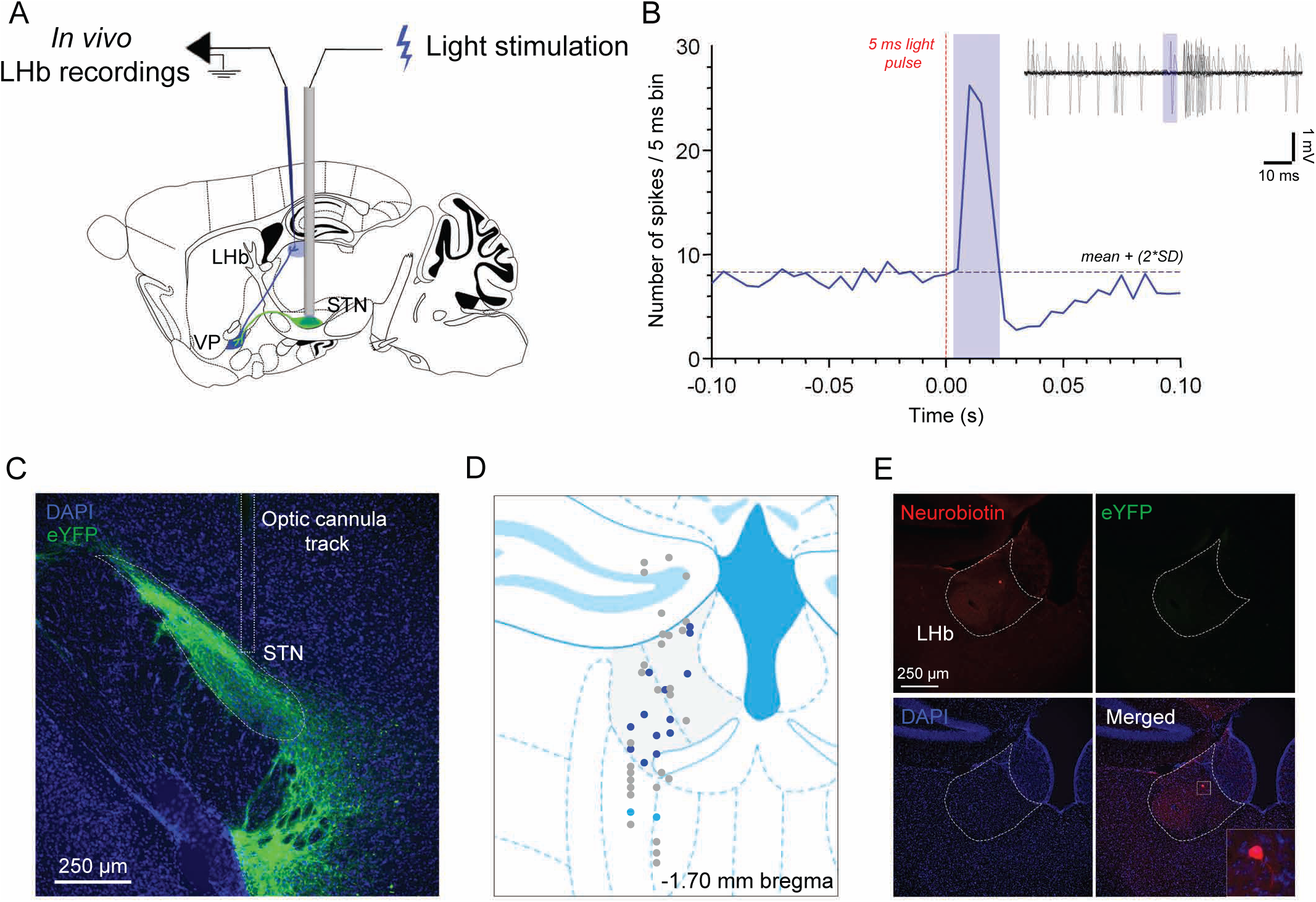
STN-photoactivation in Pitx2-ChR2-eYFP mice induces excitatory post-synaptic responses in neurons of the lateral habenula (LHb) (A) Procedure of LHb recordings experiments with STN-photostimulation in Pitx2/ChR2-eYFP mice. (B) Close-up of the averaged PSTH of LHb-excited neurons upon STN photostimulation and overlap of 10 traces of a LHb neuron centred on the 5 ms light pulse. (C) Representative image of coronal section showing eYFP fluorescence (green) in STN cell bodies of Pitx2/ChR2/eYFP mice with optic cannula track above the STN; nuclear marker (DAPI, blue); scale, 250 µm. (D) Reconstructed mapping of LHb recorded neurons (N=44 neurons recorded in 3 mice). Dark blue dots indicate LHb neurons excited during the stimulation protocol; light blue dots correspond to excited neurons outside of the LHb; grey dots correspond to neurons which did not respond to the light. (E) Excited LHb neuron injected with neurobiotin. No eYFP-positive fibers in the LHb.

### Selected optogenetic activation STN-VP projections results in VP firing

Several projections to the LHb are involved in aversion. With the lack of direct projection from the STN to LHb, we decided to assess the putative role of the VP as the bridge between the STN and LHb in the observed STN-driven aversion. While the STN projects to several basal ganglia structures, the ventral pallidal target area, VP, was of particular interest as it sends projections to both the VTA and LHb (Root et al., 2015; Tripathi et al., 2013). Further, the VP was recently proposed to play a role in integrating rewarding and aversive information (Creed et al., 2016; Itoga et al., 2016). Of mixed neurotransmitter phenotype, specifically the glutamatergic neurons of the VP were recently identified as sending direct projections to the LHb (Faget et al., 2018; Tooley et al., 2018). In addition, it has been shown that these VP neurons receive input from the STN, among several regions (Tooley et al., 2018).

Focusing on the VP, to address the effect of STN optogenetic activation on the excitability of VP neurons, sagittal slices from Pitx2/ChR2-eYFP and control mice were prepared for whole-cell patch-clamp recordings. STN inputs were stimulated with 473 nm laser pulses through an optic fiber placed above the VP (Figure 5A). VP neurons were all recorded in the area just below the anterior commissure, a region rich in ChR2-eYFP-expressing STN axon terminals (Figure 5B-C). The impact of the activation of the STN-VP pathway was first assessed in cell-attached configuration. Trains of stimulation at 20 Hz induced a strong increase in the firing activity of all recorded VP compared to basal firing activity (Figure 5D; p<0.05), suggesting a powerful excitatory drive from the STN on VP neurons. Next, STN-VP synaptic transmission was recorded in whole-cell voltage-clamp configuration which confirmed the glutamatergic nature of this pathway (Figure 5E). Altogether, these data support the strong impact of the STN-VP pathway on the excitability of VP neurons.

**Figure 5.**
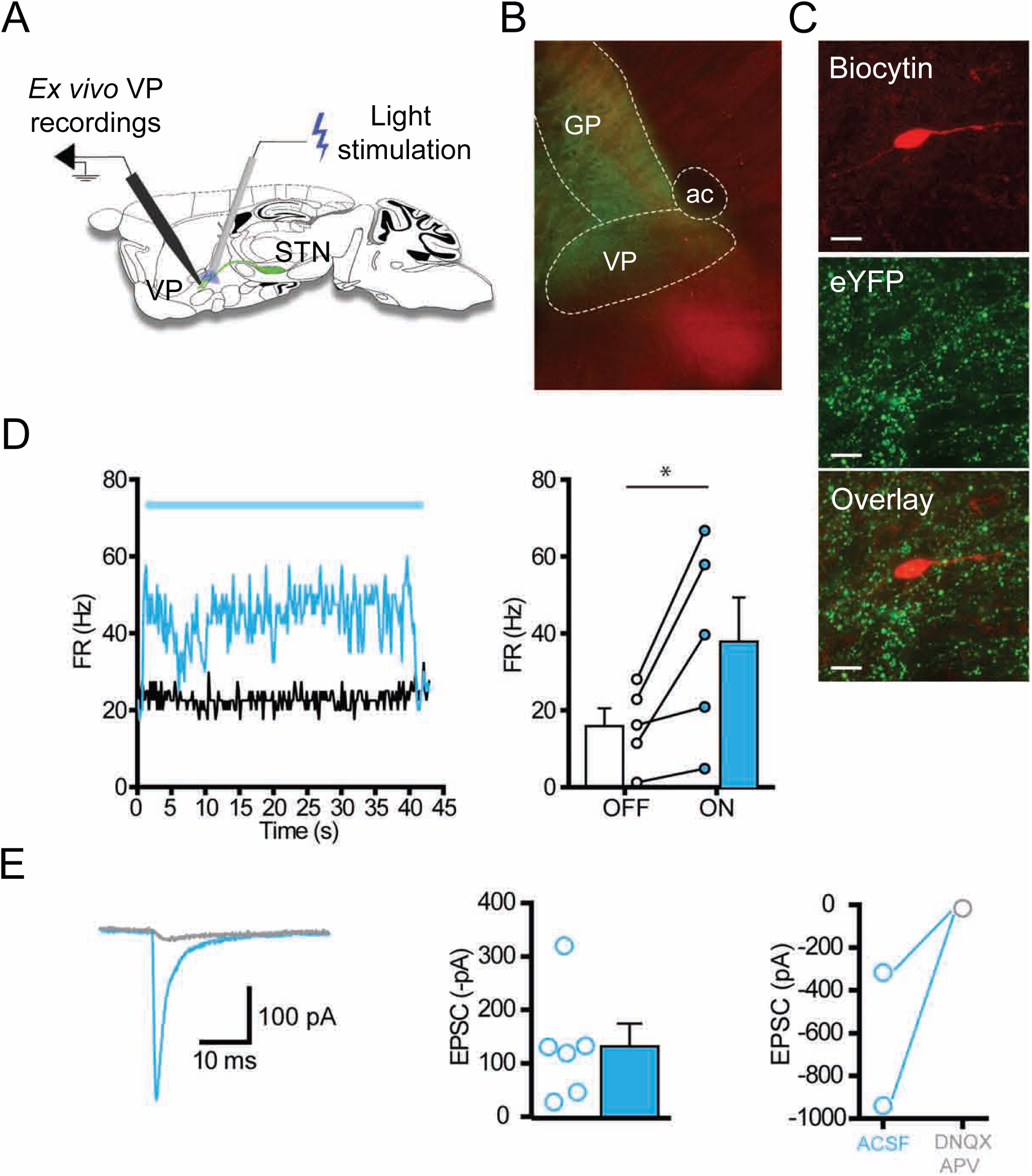
Selective optogenetic activation of projections from the STN to the ventral pallidum (VP) results in firing of VP neurons. (A) Graphical representation of a para-sagittal slice obtained from a Pitx2/ChR2-eYFP/VP mouse with a patch-clamp electrode and optic fiber placed above the ventral pallidum (VP). Epifluorescence image depicting the globus pallidus (GP) and VP. VP neurons were recorded below the anterior commissure (ac). (C) Confocal images showing a VP neuron in a Pitx2/ChR2-eYFP/VP mouse filled with biocytin (BC); surrounded by STN axon terminals expressing ChR2-eYFP (eYFP). Scale bar: 10µm. (D) Representative firing rate of a VP neuron during ON/OFF periods of light stimulation (20Hz, 1ms, for 40 s) and corresponding population data (n=5, *p<0.05, paired t-test). Note the strong increase of firing produced by light stimulation of STN axon terminals. (E) Typical light-evoked excitatory synaptic current (EPSC) which is abolished by AMPA/NMDA receptors antagonists (DNXQ, 20µm; D-APV, 50µm). (F) Population graph of light-evoked EPSCs (n=6). (F) Graph showing the pharmacological blockade of light-evoked EPSC (n=2) by DNQX and D-APV.

### Activation of STN-VP terminals induces place avoidance

Finally, to investigate if selective activation of the STN-VP pathway can drive aversive behavior, optic cannulas were bilaterally implanted above the VP in a new batch of Pitx2-Cre mice virally injected into the STN (following the terminology used above, these mice are referred to as Pitx2/ChR2-eYFP/VP and Pitx2/eYFP/VP, respectively) (Figure 6A). This allowed behavioral testing in the same RT-PP and EPM paradigms as used for the STN-photostimulation, but now upon photostimulation directed to the ChR2-eYFP-positive STN-terminals in the VP. Pitx2/ChR2-eYFP/VP mice showed an active avoidance behavior to VP-photostimulation by spending significantly less time in the light-paired compartment (A) than the unpaired compartment (B) both during the two days of real-time exposure to photostimulation (Days 3, 4), and also during the first conditioned test days (Days 5) (Figure 6B). Control mice showed an absence of stimulation-dependent response throughout the trials (Figure 6B,C).

**Figure 6.**
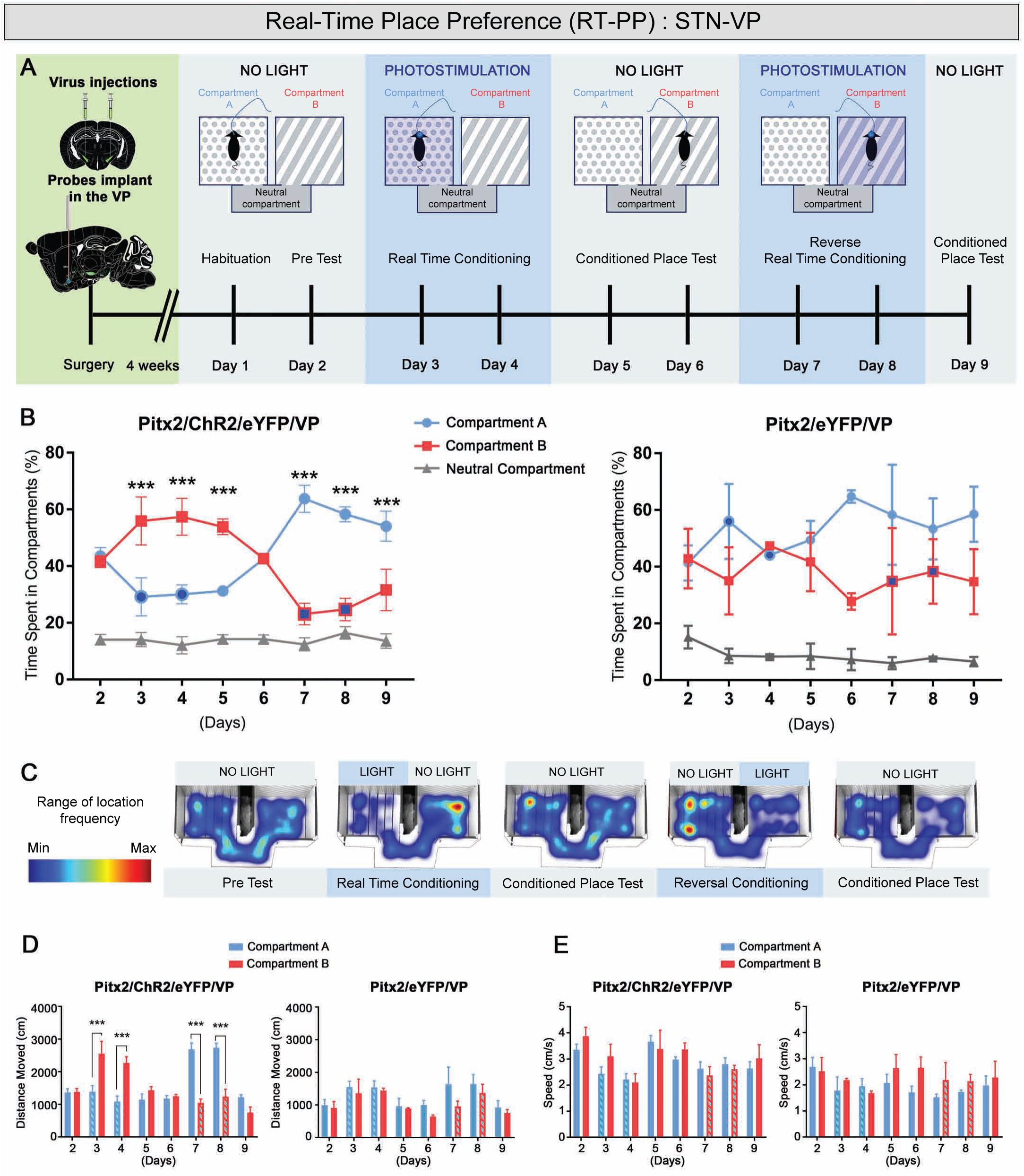
Selective photoactivation within the ventral pallidum (VP) of STN-VP projections is sufficient to drive place aversion. (A) Graphical representation of the real time place preference (RT-PP) experiment in which Pitx2/ChR2-eYFP/VP and corresponding control mice were tested. (B) Percentage of time spent in each compartment express as mean ± SEM. Dark blue filled circles (compartment A) and dark blue filled squares (compartment B) indicate photostimulation in that compartment. (Left panel) n=4 Pitx2/ChR2/eYFP/VP, compartment A vs. compartment B, ***p<0.001; (Right panel) n=3 Pitx2/eYFP/VP controls. (C) Representative occupancy heat-maps for Pitx2/ChR2-eYFP mice less time spent in the photostimulation-paired chamber than in the unpaired chamber. (D) Distance moved in each compartment expressed as mean ± SEM throughout the course of the experiment. Light blue filled histograms indicate photostimulation in that compartment. (Left panel) n=4 Pitx2/ChR2/eYFP/VP mice, compartment A vs. compartment B,***p<0.001. (Right panel) n=3 Pitx2/eYFP/VP control mice. (E) Speed in each compartment expressed as mean ± SEM. (Left panel) n=4 Pitx2/ChR2/eYFP/VP mice. (Right panel) n=3 Pitx2/eYFP/VP control mice.

Upon reversal of light-pairing (Days 7, 8), Pitx2/ChR2-eYFP/VP mice reversed their preference by again spending significantly more time in the light-unpaired compartment (A). The preference for the unpaired compartment was maintained on the final conditioned test day (Day 9) when no light was applied (Figure 6B,C). Avoidance towards the light-paired compartment was confirmed by shorter distance moved compared to the non-light paired compartment across all test days similarly to what observed upon STN optogenetic stimulation while, in the other hand, speed was not affected (Figure 6D,E).

These results suggest that the STN-VP pathway is involved in the aversive response observed upon optogenetic STN activation.

### Activation of STN-VP terminals reduces the natural preference for a sheltered environment

Also the EPM protocols were used to analyse the Pitx2/ChR2-eYFP/VP mice (Figure 7A). A similar response was observed upon VP-stimulation as was seen using STN-stimulation. That is, in the regular EPM, photoactivation of the terminals in the VP did not induce any visible difference between Pitx2/ChR2-eYFP/VP and control mice. Pitx2/ChR2-eYFP/VP mice spent most of the time in the closed arms compared to open arms and center (Figure 7B). The number of entries into either the open or closed arms were not significantly different between Pitx2/ChR2-eYFP/VP and control mice (Figure 7C). These results were in accordance with those observed with STN activation.

**Figure 7.**
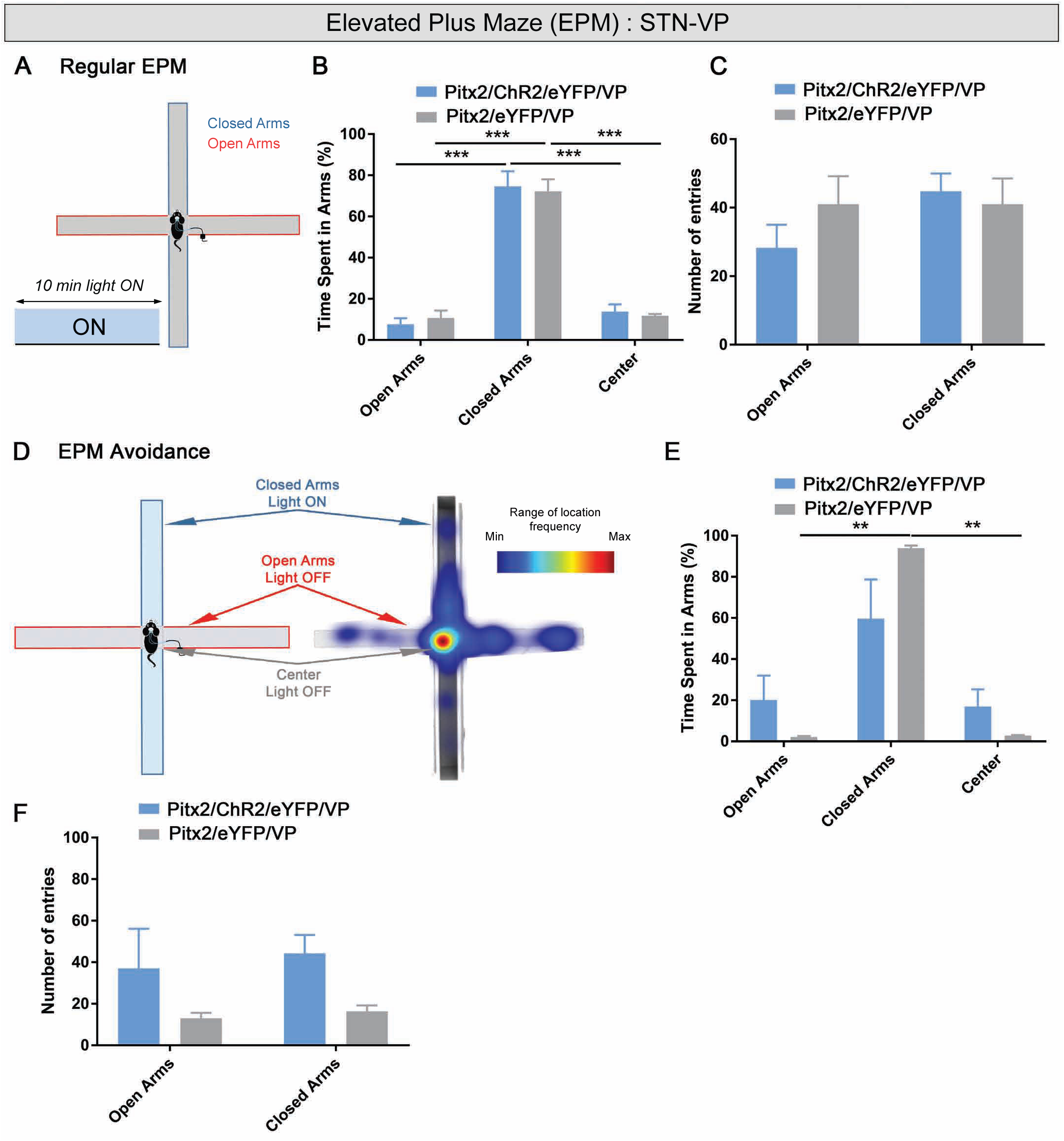
Optogenetic activation within the ventral pallidum (VP) of STN-VP terminals reduces the natural preference for a sheltered environment. (A) Schematic representation of elevated plus maze (EPM) with photostimulation delivered for 10 minutes in the entire arena. (B) Percentage of time spent in arms expressed as mean ± SEM, n=4 Pitx2/ChR2/eYFP/VP mice, open arms vs. closed arms, ***p<0.001; center vs. closed arms, ***p<0.001; n=3 Pitx2/eYFP/VP control mice, open arms vs. closed arms, ***p<0.001; center vs. closed arms, ***p<0.001. (C) Number of entries in EPM arms expressed as mean ±SEM. (D) Graphical representation of a modified version of the EPM test and representative occupancy heat map. (E) Time spent in EPM arms expressed as mean ±SEM, n=3 Pitx2/eYFP/VP, open arms vs. closed arms, **p<0.01; center vs. closed arms, **p<0.01. (F) Number of entries in arms expressed as mean ±SEM.

Next, we tested the Pitx2/ChR2-eYFP/VP mice in the re-designed EPM-avoidance paradigm (Figure 7D), where VP-photostimulation was activated directly upon entry into one of the two closed arms and disabled by leaving them. As observed upon STN-photostimulation, activation of STN-VP terminals in the VP induced a visible reduction of the time spent in the closed arms with an increase of the time spent in the open arms and center. Pitx2/eYFP/VP control mice spent significant more time in the closed arms compared to open arms and center, while Pitx2/ChR2-eYFP/VP mice spent a similar amount of time in the three compartments, annulling the natural preference for the closed arms (Figure 7E). Despite an increased tendency in the number of entries comparing Pitx2/ChR2-eYFP/VP with Pitx2/eYFP/VP control mice, no significant difference was detected (Figure 7F). In summary, while mice did not show an anxiogenic phenotype, optogenetic stimulation of STN terminals in the VP was sufficient to drive avoidance behavior.

## Discussion

Three distinct sets of optogenetics-based experiments in mice expressing ChR2 selectively in the STN enabled the identification of a previously unknown role of this clinically relevant brain area in aversion: First, we identify that photostimulation of the STN drives aversive behavior. Second, STN-photostimulation drives an excitatory response in LHb neurons. Third, photostimulation of STN-terminals in the VP results in significant firing, and induces a similar aversive response as when the STN itself is stimulated. Further, as no direct projection is observed between the STN and LHb, and with a delay in the observed evoked-firing of the LHb, the current results support an indirect connection between the STN and LHb via the VP. The identification of a causal role of the STN in driving aversion opens up for new possibilities in the current understanding and on-going decoding of aversive behavior.

The role of selective STN excitation in mediating emotional valence and motivated behaviors has so far been poorly investigated. Aversive stimuli, such as the threat of a predator or the perception of an imminent dangerous situation induce the activation of a “survival state” designed to avoid or reduce the possible harmful outcome. In this context, aversive learning allows the animals to detect the aversive stimulus and learn to actively avoid it, while aversive Pavlovian conditioning serves to associate neutral stimuli to the hostile situation and environment reducing the probability of a related behavior being expressed (Jean-Richard-Dit-Bressel et al., 2018). Obtaining a reward or avoiding punishment shapes decision-making and motivates learning with several brain regions playing a part is such vital responses (Jean-Richard-Dit-Bressel et al., 2018). Using a recently validated optogenetic approach in Pitx2-Cre mice in which STN neurons can be selectively excited and thus activated, our findings now identify aversive responses upon STN activation, possibly by acting as an upstream component of the LHb. Indeed, STN excitation caused an evoked excitation of LHb neurons, demonstrating a common circuitry, but also found the association indirect. With the observed STN-mediated excitation of VP neurons followed through with behavioral aversion, the VP is here suggested as a brain area connecting the STN and LHb. However, given the multitude of structures that communicate with the STN, additional experiments will be required to fully pinpoint the STN circuitry of aversion. *Pitx2* gene expression is characteristic of the STN, with expression starting early in STN development and persisting throughout life (Dumas and Wallén-Mackenzie, 2019; Martin et al., 2004; Schweizer et al., 2016; Skidmore et al., 2008). In previous work, we could verify the selectivity of the Pitx2-Cre transgene in driving floxed opsin constructs to the STN, with photostimulation-induced post-synaptic currents observed in pallidal (globus pallidus) and nigral (SNr) target areas (Viereckel, 2018, Schweizer 2014). Thus, in addition to the post-synaptic response of VP neurons identified here, also other STN target areas might be involved in mediating the observed behavioral avoidance. While further investigations will be needed to fully outline the neurocircuitry of STN-mediated aversion, the current findings clearly show that STN excitation caused both immediate place avoidance and cue-induced avoidance behavior, thus directly linking STN activation to aversion-processing. In this context, the STN, which in turn receives inputs from cortical regions supporting hedonic processing (Degos et al., 2008; Frankle et al., 2006; Haynes and Haber, 2013; Isaacs et al., 2018), might play a modulatory role in decision-making in response to aversive conditions.

In a recent study, we could experimentally confirm the long-assumed role of the STN in locomotion by demonstrating directly opposite motor effects by optogenetic excitation and inhibition of the STN (Guillaumin et al., 2020). We found that STN excitation was generally correlated with significant reduction in locomotor activity, while STN inhibition enhanced locomotion, just as classical models of the regulatory role of the STN in basal ganglia motor loops propose. However, we also found that these correlations were not true for all types of behavior. Upon STN-photostimulation in the open field test, jumping and self-grooming behaviors were induced, not reduced. Curiously, this was only observed in a non-challenging environment, not when mice where engaged in complex motor tasks. While jumping and grooming are distinctly measurable motor output, they might represent an emotional manifestation (Smolinsky et al., 2009; Wahl et al., 2008). In fact, the mice attempted to jump out of the experimental test arena as if to escape the stimulation. In a follow-up pilot study, we observed the mice in their natural home-cage environment. STN-photostimulation was turned on by remote control, and only in short periods. Immediately upon STN-stimulation, Pitx2/ChR2-eYFP mice responded with self-grooming and attempts to escape the home-cage. Self-grooming was mostly observed when the lid was on, preventing escape, while in an open cage, most of the mice escaped by jumping over the walls (unpublished observations).

These behavioral manifestations suggested to us that the mice experience STN activation as strongly unpleasant. Identifying a causal role for the STN in neurocircuitry of aversion could help the understanding of aversive conditions, including some of the adverse side-effects observed upon STN-DBS. In the present study, we decided to fully explore these striking observations. By implementing the RT-PP paradigm along with two different setups of the EPM, the current results now allow both the pin-pointing of place aversion upon STN activation, and also the segregation between aversion and anxiety. Further, in these experimental arenas, mice move freely between compartments in which they receive, or do not receive, optogenetic stimulation. As the mouse chooses where it will position itself throughout the test session, there is no driving force for escape. However, our current analyses clearly show that when allowed this choice, mice actively avoid areas in which they receive stimulation of either the STN itself or STN-terminals within the VP. While mice do not show measurable anxiety, significant place avoidance was evident in both the RT-PP and the EPM-avoidance tests, and this avoidance behavior was strong enough to compete out the natural aversion for an open unsheltered space.

STN activation has not been experimentally associated with aversion before. Instead, a main brain area for aversion is the LHb. During the past years, several different glutamatergic afferents to the LHb have proven responsible for mediating aversive behavior. For instance, optogenetic stimulation of glutamatergic neurons of the EP (Shabel et al., 2012; Stephenson-Jones et al., 2016), VP (Faget et al., 2018; Tooley et al., 2018) and lateral hypothalamus (Lazaridis et al., 2019; Lecca et al., 2017) projecting to the LHb all induce place avoidance. Now, our findings identifying optogenetically induced aversion in the RT-PP and EPM avoidance tests upon stimulation of either the STN itself or its terminals in the VP, place the STN as a distinct brain nucleus responsible for the control of aversion by the recruitment of VP neurons. Using *in vivo* extracellular recordings, we confirmed the activational response of the STN neurons themselves upon photostimulation, and also demonstrated that stimulation of STN glutamatergic neurons drive neuronal activity in the LHb, while patch-clamp results identify excitatory post-synaptic response in the VP. Together, these electrophysiological and behavioral findings provide strong support for an STN-VP-LHb pathway for aversion. Our findings are in accordance with recent observations identifying that activation of glutamatergic projection from several regions to the LHb triggers place avoidance as well as activation of LHb activity. Further, our results allow the identification of one more component to the aversion circuitry, and establish the STN as putative upstream regulator of the VP-LHb pathway.

Curiously, the VP is mostly a GABAergic structure, but recent studies demonstrated that it also contains glutamatergic neurons projecting to the LHb (Faget et al., 2018; Tooley et al., 2018). The glutamatergic population of the VP was recently shown to receive projections from the STN, among several areas, and selective stimulation of glutamatergic VP neurons was demonstrated to evoke EPSCs in the LHb and induce place avoidance (Tooley et al., 2018). The glutamatergic population of the VP receives input from different cortical and midbrain regions (Knowland et al., 2017; Tooley et al., 2018), and it remains to establish which of these upstream structures, if any, is responsible for their activation in avoidance behavior. However, our present results pin-point the STN as one distinct subcortical regulator of VP activity. Further, the findings identify that the multisynaptic STN-VP-LHb circuit is composed mainly of glutamatergic connections, which provides new insight into the role of excitatory pathways in aversive behavior.

The clinical importance of the STN highlights the need for decoding the role of this small nucleus in behavioral regulation. STN-DBS treatment for motor dysfunction in advanced stage Parkinson’s disease is a long established and highly successful clinical intervention (Benabid et al., 2009). More recently, STN-DBS has also been approved as treatment for highly refractory and treatment-resistant OCD, another disorder in which the STN shows aberrant activity (Klavir et al., 2009; Mallet et al., 2008; Winter et al., 2008). However, considering that numerous adverse side-effects of STN-DBS have been reported, including low mood state, depression, personality changes and even suicide (Péron et al., 2013; Temel et al., 2006), manipulating the STN might come with a certain risk. While this issue has been challenging to resolve, any direct causality between STN excitation and behavioral aversion in mice is clearly crucial to take into consideration as putatively important not only experimentally, but also clinically.

The present results underscore the importance of decoding the full repertoire of behavioral regulation executed by the STN not least to improve treatment prediction and outcome in any intervention aiming to manipulate the STN and its pathways. Beyond Parkinson and OCD, STN-DBS has recently gained attention as treatment for additional neuropsychiatric disorders, including addiction (Pelloux and Baunez, 2013). This interest is strongly based on findings that STN-DBS reduces symptoms of the Dopamine dysregulation syndrome in Parkinson’s disease (Bandini et al., 2007; Broen et al., 2011; Knobel et al., 2008; Smeding et al., 2006). Also studies in rodents showing that experimental STN-DBS induces different motivational responses depending on the nature of the reinforcers (Baunez et al., 2002; Darbaky et al., 2005; Rouaud et al., 2010), and can reduce addictive behaviors (Lardeux and Baunez, 2008; Rouaud et al., 2010; Wade et al., 2017), have high-lighted the STN in addiction. However, it has also been shown that STN lesioning in rats led to altered emotional state in response to various rewarding and aversive stimuli (Pelloux et al., 2014). Taken together, both experimental and clinical data high-light the STN structure as important in affective processing.

Here, we describe the identification of the STN as a non-canonical source of negative emotional value. Well embedded within the brain circuitry of negative reinforcing properties, the results point towards a pivotal role of the clinically relevant STN in aversive learning. Evidently, aberrant activity of an STN-VP-LHb circuitry, as well as its manipulation, could result in both beneficial and detrimental effects. Further studies will be needed to pinpoint the nature of all pathways emerging from the STN that have an impact on rewarding *vs* aversive responses. In a recent transcriptomics-histological effort, we identified gene expression patterns that define distinct spatio-molecular domains within the STN structure (Wallén-Mackenzie et al., 2020). Based on the current finding that STN encodes aversive behavior, future experimental approaches should make an effort to correlate specific domains within the STN with behavioral output. Such anatomical-functional mapping within the STN structure would be important not least to aid towards improving spatial selectivity in STN-based interventions.

## METHODS SECTION

### Animal housing and ethical permits

Pitx2-Cre_C57BL/6NTac transgenic mice were bred in-house and housed at the animal facility of Uppsala University before and during behavioral experiments or at University of Bordeaux after virus injections for *in vivo* electrophysiological experiments. Mice had access to food and water *ad libitum* in standard humidity and temperature conditions and with a 12 hour dark/light cycle. PCR analyses were run to confirm the genotype of Pitx2-Cre positive mice from ear biopsies. All animal experimental procedures followed Swedish (Animal Welfare Act SFS 1998:56) and European Union Legislation (Convention ETS 123 and Directive 2010/63/EU) and were approved by local Uppsala or Bordeaux (N°50120205-A) Ethical Committee.

### Stereotaxic virus injection and optic cannula implantation

#### Virus injection

Stereotaxic injections were performed on anesthetized Pitx2-Cre mice maintained at 1.4-1.8 % (0.5-2 L/min, isoflurane-air mix v/v). Before starting the surgery, and 24h post-surgery, mice received subcutaneous injection of Carprofen (5 mg/Kg, Norocarp). A topical analgesic, Marcain (1.5 mg/kg; AstraZeneca), was locally injected on the site of the incision. After exposing the skull, holes were drilled in the skull for virus injections and skull screws implantations for behavioral experiments. Mice were injected in the STN bilaterally with a virus containing either a Cre-dependent Channelrhodopsin (rAAV2/EF1a-DIO-hChR2(H134R)-eYFP) named Pitx2/ChR2-eYFP, or an empty virus carrying only a fluorescent protein (rAAV2/EF1a-DIO-eYFP) named Pitx2/eYFP, respectively 3.8×10^12^ virus molecules/mL, 2.7×10^12^ virus molecules/mL; 4.6×10^12^ virus molecules/mL (viruses purchased from UNC Vector Core, Chapel Hill, NC, USA) at the following mouse brain coordinates (from Paxinos and Franklin, 2013): anteroposterior (AP) = −1.90 mm, mediolateral (ML) = +/− 1.70 mm from the midline and 250 nL of virus was injected with a NanoFil syringe (World Precision Instruments, Sarasota, FL, USA) at two dorsoventral levels (DV) = −4.65 mm and − 4.25 mm from the dura matter at 100 nL.min-1.

#### Optic cannula implantation

Optic cannulas (Doric Lenses) were implanted directly after completion of virus injections. Two skull screws were implanted in the skull to hold the optic cannula-cement-skull complex. Primers were then applied and harden with UV light (Optibond). Optic cannulas were implanted bilaterally above the STN (coordinates: AP = − 1.90 mm, ML = +/−1.70 mm from the midline DV = −4.30 mm) or the VP (coordinates: AP = +0.45 mm, ML = +/−1.55 mm from the midline, DV = −4.00 mm), and fixed with dental cement. 1 mL of saline was injected subcutaneously at the end of the surgery.

### Single-cell extracellular recordings

#### Surgery

*In vivo* single cell extracellular recordings started at least 4 weeks after virus injections. Mice were anesthetized with a mix isoflurane-air (0.8-1.8 % v/v) and placed in a stereotaxic apparatus. Optic fiber, optrode and glass micropipettes coordinates were AP = − 1.90 mm, ML = +/− 1.70 mm and DV = −4.30 mm for the STN and AP = −1.60 mm, ML = +/− 0.50 mm, DV = −2.00 to −3.30 mm for the LHb.

#### STN Optotagging

A custom-made optrode was made with an optic fiber (100 µm diameter, Thorlabs) connected to a laser (MBL-III-473 nm-100 mW laser, CNI Lasers, Changchun, China) mounted and glued on the recording glass micropipette which was filled with 2% pontamine sky blue in 0.5 µM sodium acetate (tip diameter 1-2 µm, resistance 10-15 MΩ). The distance between the end of the optic fiber and the tip of the recording pipette varied between 650 nm and 950 nm. Extracellular action potentials were recorded and amplified with an Axoclamp-2B and filtered (300 Hz/0.5 kHz). Single extracellular spikes were collected online (CED 1401, SPIKE2; Cambridge Electronic Design). The laser power was measured before starting each experiment using a power meter (Thorlabs). The baseline was recorded for 100 seconds for each neuron before starting any light stimulation protocols which were set and triggered with Spike2 software. Light protocol consisted in a peristimulus time histogram (PSTH, 0.5 Hz, 5 ms pulse duration, 5-8 mW) for at least 100 seconds followed, after returned to baseline, by a “behavioral” protocol corresponding to the parameters used for behavioral experiments (20 Hz, 5 ms pulse duration, 5-8 mW).

#### LHb recordings

An optic fiber and a recording pipette filled with either 2% pontamine sky blue or 2% neurobiotin were respectively positioned in the STN and LHb. For each LHb neurons, a PSTH was recorded upon STN optogenetic stimulation (PSTH 0.5 Hz, 5 ms pulses, 5-8 mW) for at least 100 seconds. Neurons are considered as excited during the PSTH protocol when, following the light pulses centered on 0, the number of spikes/5ms bin is higher than the baseline (−500 ms to 0 ms) plus two times the standard deviation. Injection of neurobiotin (2% neurobiotin in 0.5 M acetate sodium) by juxtacellular iontophoresis was performed in some cases for precise spatial identification of excited neurons.

### *Ex vivo* electrophysiology

STN and VP neurons were recorded in acute brain slices, prepared as previously described (Dupuis et al., 2013; Froux et al., 2018). Briefly, Pitx2/eYFP and Pitx2/ChR2-eYFP mice (> 12 months) were deeply anesthetized with an i.p. injection of ketamine/xylazine (75/10 mg/kg) mixture and then perfused transcardially with cold (0-4°C) oxygenated (95% O_2_ - 5% CO_2_) artificial cerebrospinal fluid (ACSF), containing the following (in mM): 230 sucrose, 26 NaHCO_3_, 2.5 KCl, 1.25 NaH_2_PO_4_, 0.5 CaCl_2_, 10 MgSO_4_ and 10 glucose (pH∼7,35). The brain was quickly removed, glued to the stage of a vibratome (VT1200S; Leica Microsystems, Germany), immersed in the ice-cold ACSF and sectioned into 300 μm thick parasagittal slices. Slices were transferred for 1 hour to a standard oxygenated ACSF solution, warmed (∼35°C) containing (in mM unless otherwise stated): 126 NaCl, 26 NaHCO_3_, 2.5 KCl, 1.25 NaH_2_PO_4_, 2 CaCl_2_, 2 MgSO_4_, 10 glucose, 1 sodium pyruvate and 4.9 μML-gluthathione reduced. Single slices were then transferred to a recording chamber, perfused continuously with oxygenated ACSF (without sodium pyruvate and L-gluthathione) heated at 32-34°C, and visualized using infrared gradient contrast video microscopy (Ni-e workstation, Nikon) and a 60X water-immersion objective (Fluor APO 60X/1.00 W, Nikon).

Recordings from individual STN and VP neurons were performed with patch electrodes (impedance, 3–8 M*Ω*)obtained from borosilicate glass capillaries (GC150F10; Warner Instruments, Hamden, CT, USA) pulled with a horizontal puller (P-97; Sutter Instruments, Novato, CA, USA). For cell-attached and whole-cell voltage-clamp recordings, pipettes were filled with K-Gluconate-based internal solution containing (in mM): 135 K-gluconate, 3.8 NaCl, 1 MgCl_2_.6H_2_O, 10 HEPES, 0.1 EGTA, 0.4 Na_2_GTP, 2 Mg_1.5_ATP, 5 QX-314 and 5.4 biocytin (pH=7.2, ∼292 mOsm). Recordings were obtained using a Multiclamp 700B amplifier and Digidata 1440A digitizer controlled by Clampex 10.2 (Molecular Devices LLC). Signals were sampled at 20 kHz and low-pass filtered at 4 kHz. Series resistance was monitored throughout the experiment by voltage steps of −5 mV. Data were discarded when the series resistance changed by >20%.Biocytin-filled neurons were identified. For *ex vivo* optogenetic stimulation, a LED laser source (Prizmatix, Israel) connected to optic fiber (∅: 500 µm) was placed above the brain slice. Light intensities ranged from 4mW to 90mW at the tip of the optic fiber depending of the type experiments. For cell body stimulation, continuous 100ms long duration light stimulation was applied at low (4mW) and high (90 mW) intensities. To evoke synaptic transmission, single pulses or train of stimulation (800 pulses at 20Hz) of 1ms duration at full power (90mW) were used in order to maximize axon terminal depolarization and efficient release of neurotransmitter. After electrophysiological recordings, slices were fixed overnight in 4% paraformaldehyde, and maintained in PBS-azide at 0.2% at 4°C until immunohistochemical processing.

### Histological analysis

Following behavioral analyses, mice were deeply anesthetized and perfused transcardially with phosphate-buffer-saline (PBS) followed by ice-cold 4% formaldehyde. Brains were extracted and 60 μm sections were cut with a vibratome.

Fluorescent immunohistochemistry was performed to enhance the eYFP signal. All mice were analysed. After rinsing in PBS, sections were incubated for 90 min in PBS 0.3% X-100 Triton containing 10% blocking solution (normal donkey serum) followed by an incubation with primary antibody (chicken anti-GFP 1:1000, cat. no. ab13970, Abcam), diluted in 1% normal donkey serum in PBS, overnight at 4°C. Next day, sections were rinsed in PBS plus 0.1% Tween-20 solution and incubated for 90 min with secondary antibody diluted in PBS (A488 donkey anti-chicken 1:1000, cat. no. 703-545-155, Jackson Immunoresearch). After rinsing in PBS containing 0.1% Tween-20, sections were incubated for 30 min with DAPI diluted in distilled water (1:5000). Sections were mounted with a Fluoromount aqueous mounting medium (Sigma, USA) and cover-slipped. Sections were scanned with NanoZoomer 2-0-HT.0 (Hamamatsu) scanner and visualized with NDPView2 software (Hamamatsu).

For *in vivo* electrophysiological experiments, a deposit was made at the last recording coordinates by pontamine sky blue iontophoresis (−20 μA, 25 min). Brains were then collected after 0.9% NaCl and 4% PFA transcardial perfusion. 60 µm brain sections were cut with a vibratome at the levels of the STN and the LHb, mounted with Vectashield medium (Vector Laboratories), coverslipped and visualized with an epifluorescent microscope to confirm eYFP expression, recording pipette and optic fiber positions. Neurons injected with neurobiotin were revealed by first incubating the brain slices in a solution of saturation/permeabilization (1X PBS-Triton X-100 0.3% and 1% BSA) for one hour at room temperature followed by an incubation overnight at room temperature in a solution containing 1X PBS-Triton X-100 0.3% (v/v; Sigma-Aldrich) and Streptavidine-557 (1:500; R&D System #NL 999).

To reveal biocytin-filled neurons after *ex vivo* recordings, brain slices were incubated overnight in a solution of PBS 0.01 mM/Triton™ X100 0.3% (v/v; Sigma-Aldrich) containing streptavidine-557 (1/1000; Cat# NL999; R&D system). Fluorescent immunochemistry was used to enhance eYFP signal as described above but with a mouse anti-GFP (1/600; Cat#11814460001, Roche or Cat#A11122; Life technologies) primary antibody and an Alexa-Fluor Donkey anti-Mouse 488 (1/500; Cat#A21202, Life technologies) as secondary antibody. The slices were then washed in PBS 0.01 mM before mounting onto slides in Vectashield medium (CAT#H-1000, Vector laboratories). Images were acquired with a fluorescence microscope (Axio Imager 2, Zeiss) or a confocal microscope (Leica, SP8) and were analysed with Fidji.

### Behavioral testing

After approximately four weeks of recovering from surgery, mice were tested in validated behavioral tests by an experimenter blind to experimental groups, during which the same stimulation protocol was used for Pitx2/ChR2-eYFP, Pitx2/ChR2-eYFP/VP and respective controls: 473 nm light, 5 mW, 20 Hz, 5 ms pulse delivered by a MBL-III-473 nm-100 mW laser (CNI Lasers, Changchun, China). Duration and condition of stimulation are specified for each test. After completed behavioral tests, mice were sacrificed and brains analysed histologically and for assessment of optic cannula position. Mice in which the optic cannulas were not in correct position were excluded from the analysis. For each test, results were obtained from groups of mice that followed the same battery of experiments. The initial sample size was determined based by previous published studies with similar experimental design.

### Habituation

Three weeks after surgery and before the first behavioral test, all mice were handled and habituated to the experimental room and to the optic cables to reduce the stress during the day of the experiment. Before each behavioral test, mice were acclimatized for 30 minutes in the experimental room.

### Real-time place preference (RT-PP)

A three-compartment apparatus (Spatial Place Preference Box, Panlab, Harvard Apparatus) was used for the RT-PP test. The apparatus is composed of two compartments with different visual and tactile cues for both floors and walls and a connecting corridor (neutral compartment) with transparent walls and floor. The study was carried out throughout 8 consecutive days preceded by one *“Habituation”* day. On each day of the experiment, subjects were placed in a neutral cage for at least 3 minutes to recover after connecting the optic cables. Mice were subsequently moved into the transparent corridor of the apparatus where, after 30-50 s, the doors were removed and animals allowed to freely explore the apparatus. The position of the mouse was detected by a camera positioned above the apparatus. On *“Habituation”, “Pre Test”* and “*Conditioning Place Test”* days (15 min), mice were connected to the optic fiber but no light was paired to any chamber. On the “*Real Time Conditioning”* days (30 min), one chamber was randomly chosen as a light-paired chamber (counterbalanced across animals) while the other one was subsequently assigned as light-paired chamber on the “*Reverse Real Time Conditioning”* days (30 min). Every time the subject entered the light-paired chamber, the laser was activated according to the stimulation protocol. Cumulative duration of the time spent in each compartment was recorded by using the Ethovision XT13.0/14.0 tracking software (Noldus Information Technology, The Netherlands). Mice that showed strong initial preference during the “Pre Test” for either one of the two compartments (<25% or >75% of time spent) were excluded from the statistical analysis.

### Elevated plus maze (EPM)

EPM analysis was carried out in an apparatus with the shape of a plus, made of two open arms (35 cm length) and two closed arms (35 cm length) with walls (15 cm high) and crossed in the middle to create a center platform. The EPM apparatus adjusted for mice is elevated 50 cm from the floor. Mice were placed in a neutral cage for 5 minutes to recover from optic cable connection. Mice were next placed individually in the center of the maze facing one of the open arms and allowed to freely explore the apparatus for 10 minutes with light delivered for the total duration of the test according to the stimulation protocol. The results of the test were recorded with a camera placed above the EMP arena. Time spent in arms and number of entries in arms were manually scored by experimenter blind to experimental groups through subsequent video analysis.

### Modified version of the EPM (EPM avoidance)

The re-designed version of the EPM test aims to assess the aversive effect induced by the photostimulation in comparison to the natural aversion experienced in the open arms of the apparatus. In the EPM avoidance test, photostimulation was activated upon entry into any of the closed arms, and disabled by leaving it. In contrast, visiting the open arms or occupying the center zone had no effect on photostimulation. The animal was placed in the center zone of the arena and allowed to freely explore the apparatus for 15 min. Time spent in each compartment and number of entries were recorded by the Ethovision XT13.0/14.0 tracking software (Noldus Information Technology, The Netherlands).

### Statistics

Data are expressed on the plots as means ± SEM. For behavioral analysis, repeated measures two-way ANOVA were performed followed by Bonferroni’s or Tukey’s multiple comparisons where appropriate. For single cell extracellular recordings, Friedman test was performed followed by Dunn’s multiple comparisons. For *ex vivo* electrophysiology, paired t-test was used (GraphPad Prism version 7.00 for Windows, GraphPad Software, La Jolla California USA). See Supplementary file 1.

---

**Supplementary file 1**

Detailed description of the statistics used for the different experiments.

## Acknowledgements

We thank Prof James Martin, Baylor College of Medicine, Houston, Texas, USA, for generously providing the Pitx2-Cre transgenic mouse line. Members of the Mackenzie laboratory are thanked for feedback throughout the study. This work was supported by Uppsala University and by grants to Å.W.M from the Swedish Research Council (SMRC 2017-02039), the Swedish Brain Foundation (Hjärnfonden), Parkinsonfonden, and the Research Foundations of Bertil Hållsten, Zoologiska and Åhlén.

